# Isolation and characterization of key genes that promote flavonoid accumulation in purple-leaf tea (*Camellia sinensis* L.)

**DOI:** 10.1101/139105

**Authors:** Xiujuan He, Xuecheng Zhao, Liping Gao, Xingxing Shi, Xinlong Dai, Yajun Liu, Tao Xia, Yunsheng Wang

## Abstract

There were several high concentrations of flavonoid components in tea leaves that present health benefits. A novel purple-leaf tea variety, ‘Mooma1’, was obtained from the natural hybrid population of Longjing 43 variety. The buds and young leaves of ‘Mooma1’ were displayed in bright red. HPLC and LC-MS analysis showed that anthocyanins and O-Glycosylated flavonols were remarkably accumulated in the leaves of ‘Mooma1’, while the total amount of catechins in purple-leaf leaves was slightly decreased compared with the control. A R2R3-MYB transcription factor (*CsMYB6A*) and a novel UGT gene (*CsUGT72AM1*), that were highly expressed in purple leaf were isolated and identified by transcriptome sequencing. The over-expression of transgenic tobacco confirmed that *CsMYB6A* can activate the expression of flavonoid-related structural genes, especially *CHS and 3GT,* controlling the accumulation of anthocyanins in the leaf of transgenic tobacco. Enzymatic assays *in vitro* confirmed that CsUGT72AM1 has catalytic activity as a flavonol 3-O-glucosyltransferase, and displayed broad substrate specificity. The results were useful for further elucidating the molecular mechanisms of the flavonoid metabolic fluxes in the tea plant.

## 1. Introduction

Flavonoids, produced from the phenylpropanoid pathway, are a large group of plant secondary compounds, which fulfill important functions such as defending pathogen infection (Jasinski *et al*., 2009), avoiding the damage from UV irradiation (Li *et al*., 1993), and interaction of the plant with temperature and other environments (Azuma *et al*., 2012; Cohen *et al*., 2012). There are mainly six groups of flavonoids in plant tissues, which include flavan-3-ols (catechins and proanthocyanidins), anthocyanins, flavanonols, flavonols, flavones and phenolic acid (Winkel-Shirley, 2001). The key components of flavonoids and anthocyanins are responsible for the attractive colors seen in various plant organs (such as leaf, flower, and fruit), avoiding excess light damage, and aiding pollination and seed dispersal (van Tunen and Mol, 1991). In recent years, the health benefit of anthocyanins received more significant attention. Scientists have systematically investigated and confirmed that anthocyanins could reduce the risk of lifestyle-related diseases, such as hypertension (Jennings *et al*., 2012), liver disorder (Chang *et al*., 2013), cerebral disorder (Rahman *et al*., 2008), dysentery and diarrhea (Hidalgo *et al*., 2012), and urinary problems (Kuo *et al*., 2012).

Tea (*Camellia sinensis*) is the most widely consumed non-alcoholic beverage. It constitutes rich catechins (flavan-3-ol) flavonoids, and around 12–24% of the dry mass of the younger leaves (Ho *et al*., 2009). Nevertheless, the concentration of anthocyanins is few to zero in normal tea leaves. Recently, researchers have developed special purple-leaf tea varieties in different tea growing countries (Joshi *et al*., 2015; Lv *et al*., 2015; Saito *et al*., 2011; Terahara *et al*., 2001). These special tea varieties have been shown to contain high quantity of anthocyanins. There are eight anthocyanins that are isolated and identified from anthocyanin-rich tea (Saito *et al*., 2011). The health benefits of purple-leaf tea have been preliminarily studied (Hsu *et al*., 2012; Maeda-Yamamoto *et al*., 2012). Hsu et al study confirmed that the anthocyanin-rich tea is considered as a novel dietary compound for colorectal cancer chemoprevention (Hsu *et al*., 2012). However, there was no major systematic or global analysis reported on the molecular mechanism of anthocyanin accumulation in these tea varieties.

Anthocyanins and other flavonoid compounds are derived from the phenylpropanoid and flavonoid biosynthetic pathways. In recent years, most of the functional and regulatory genes involved in these flavonoid pathways are evident through model species. In this pathway, the functional enzymes have been classified into two groups, early-biosynthetic enzymes are required for the synthesis of flavonoids, and late-biosynthetic enzymes are used for the synthesis of anthocyanins, catechins, and flavonols (Dixon *et al*., 2005; Tanner *et al*., 2003; Winkel-Shirley, 1999; Winkel, 2006). UDP-glycosyltransferase (UGT) is the last enzyme produced during the anthocyanin and flavonol biosynthesis. It helps to catalyze the transfer of glucosyl moiety from UDP-glucose to produce the first stable pigment (Chen *et al*., 2011). However, because of the competition in the biosynthetic pathways for substrate and multiplex branches, the relationship among plant flavonoids still remain unclear.

Several studies have reported the polymorphic characteristics of some key genes, which in turn determined the fluxion of flavonoid pathway(Nesi *et al*., 2001). Numerous studies have suggested that the fluxion of this pathway is tightly regulated by the transcription factor complex MYB-bHLH-WD40 (MBW). MBW complexes are also known to control various aspects of epidermal cell patterning, such as the development of trichomes and root hairs(Wang *et al*., 2010). Specific combinations of R2R3-MYB transcription factor with bHLH and WD40 regulate specific pathways of anthocyanin or PA biosynthesis (Terrier *et al*., 2009; Winkel-Shirley, 2001). The accumulation of anthocyanins requires the activity of MBW complex consisting of MYB (including PAP1, PAP2, MYB113, MYB114), TT8, and TTG1 in *Arabidopsis* (Gonzalez *et al*., 2008). Some studies have indicated that R2R3-MYB transcription factors play a central role in distinguishing the target gene in these pathways. Zhao et al had predicted that R2R3-MYB genes were involved in the flavonoid biosynthesis of *C. sinensis* (Zhao *et al*., 2013). However, *C.sinensis* lacked Sg6 CsMYB genes, which might be due to low anthocyanin content in the tea plants.

According to the report from a latest study by Sun et al, isolation of transcription factor R2R3-MYB anthocyanin 1 (*CsAN1*) from purple-leaf tea variety ‘Zijuan’ conferred ectopic accumulation of anthocyanins in purple-leaf tea (Sun *et al*., 2016). In this study, a purple-leaf tea variety ‘Mooma 1’ was obtained from the natural hybrid population of Longjing 43 variety in Shitai, Anhui, China (latitude 30.15 N, longitude 117.50 E). Buds, young leaves and stems of ‘Mooma 1’ appear red in the spring. A R2R3-MYB factor (*CsMYB6A*), and a novel UGT gene (*CsUGT72AM1*), were isolated and identified by transcriptome sequencing. The component analysis, quantitative reverse transcriptase PCR (qRT-PCR) analysis, and the over-expression of transgenic tobacco confirmed that *CsMYB6A* can activate the expression of flavonoid-related structural genes controlling the accumulation of anthocyanins and flavonols in the leaves. Enzymatic in vitro assays of CsUGT72AM1 confirmed that CsUGT72AM1 has catalytic activity as flavonol 3-O-glucosyltransferase, and displays broad substrate specificity. The results are useful to further elucidate the molecular mechanisms of the flavonoid metabolic fluxes in the tea plant.

## 2. Materials and methods

### 2.1 Plant material and standard chemicals

Leaves of tea variety ‘Mooma 1’and wild type ‘Longjing 43’(Camellia Sinensis CV ‘Longjing’) were sampled from healthy pants that are grown in the tea garden of Shitai County, Anhui, China (latitude 30.19 N, longitude 4 E, altitude 20 m above mean sea level). A total of 10 buds and leaves were randomly collected from different branches, frozen immediately in liquid nitrogen, and stored at −80°C for RNA-seq, qRT-PCR and HPLC analysis.

Catechin (C), gallocatechin (GC), epicatechin (EC), epigallocatechin (EGC), epicatechin 3-O-gallate (ECG), epigallocatechin 3-O-gallate (EGCG) were obtained from Shanghai RongHe Pharmaceutical Co. Myricetin 3-O-glucoside, quercetin 3-O-glucoside, kaempferol, petunidin, cyanidin, cyanidin 3-O-glucoside, and delphindin were purchased from Sigma Chemicals Co.

### 2.2 Tissue slicing and staining

Samples for observation were prepared by standard free-hand sectioning (Lux *et al*., 2005). To slice the tea tissues, fresh carrot was used as a supporter (Liu *et al*., 2009). The section was observed under a microscope (Olympus, Tokyo, Japan). The sections were stained using 0.01% (w/v) 4-dimethylaminocinnamaldehyde (DMACA) in absolute ethanol containing 0.8% w/v hydrochloric acid to observe the presence of catechins in the tissue of tea leaf (Abeynayake *et al*., 2011).

### 2.3 Analysis of catechins, flavonols, and anthocyanins in leaves

Leaves were ground to a fine powder in liquid nitrogen. The powder (1 g) was extracted with 5 ml methanol at room temperature for 10 min, followed by centrifugation at 4000 × g for 15 min. The residue was re-extracted thrice by this method. The supernatants were filtered through a 0.22 μm membrane. Catechins, flavonols, and anthocyanins were analyzed according to the liquid chromatography–mass spectrometry (LC-MS) and HPLC methods (Wang *et al*., 2012). Catechins, flavonols, and anthocyanins were quantified at 280 nm, 345 nm, and 530 nm respectively.

Since only 11 standards were available, myricetin 3-O-glucoside was used as the molar equivalent to quantify its derivatives, quercetin 3-O-glucoside for all the quercetin 3-O-glycosides, kaempferol for kaempferol 3-O-glycosides, petunidin for petunidin 3-O-glycosides, cyanidin for cyanidin 3-O-glycosides, and delphindin for delphindin 3-O-glycosides. All samples were run in triplicate for both quantitation and multivariate statistical analysis.

### 2.4 cDNA library construction and transcriptome sequencing

Total RNA of tea and tobacco (*Nicotiana tabacum*) leaves was isolated with RNAiso Plus and RNAiso-mate for Plant Tissue kits and treated with DNase I according to manufacturer’s instructions (Takara, China). RNA quality was examined using 1% agarose gel and the concentration was determined using a Nanodrap spectrophotometer (Thermo, Waltham, MA, USA). Illumina sequencing was performed at Beijing Genomics Institute (BGI, Wuhan, China) on the HiSeq™ 2000 platform (Illumina, San Diego, CA). The de novo assembly, functional annotation, and metabolic pathway analysis were carried out by BGI Institute according to the manufacturer’s instructions. Genes involved in flavonoid pathway were analyzed using Camellia sinensis unigenes as illustrated in Figure 3.

### 2.5 qRT-PCR analysis

Quantitative real-time PCR (qRT-PCR) was performed using the SYBR^®^ Premix Ex Taq™ II (Perfect Real Time) kit (Takara, Japan) on a Peltier Thermal Cycler PTC200 (Bio-Rad, USA) with gene-specific primer pairs (Table S1). The related expression level was normalized against the expression level of the housekeeping gene glyceraldehyde-3-phosphate dehydrogenase (GAPDH) in tea plant (Jiang *et al*., 2013) or actin in tobacco (Pang *et al*., 2007). The melting curve was performed to determine the PCR product size and to detect possible primer dimers. Triplets of all samples were run. The cycle number at which the reaction crossed an arbitrarily placed threshold (CT) was determined for each gene, and the relative expression of each gene was determined using the equation 2^−ΔΔCT^, where ΔΔCT = (CT_Target_ – CT_GAPDH/Actin_)_sample_ - (CT_Target_ - CT_GAPDH/Actin_)_control_ (Livak and Schmittgen, 2001).

### 2.6 Transformation of tobacco plants with CsMYB6A

The Gateway Cloning System was used to construct the transformation vectors of CsMYB6A (Lei *et al*., 2007). The PCR primer pairs for linking the attB adaptors are listed in the Additional file 2: Table S1. CsMYB6A PCR product was cloned into the entry vector pDONR207 by Gateway BP Clonase Enzyme mix according to the manufacturer’s instructions (Invitrogen, USA). The pDONR207-CsMYB6A entry vector was then transferred into the Gateway plant transformation destination vector pCB2004 using Gateway LR Clonase (Invitrogen, USA). Recombinant colonies pCB2004-CsMYB6A and control pCB2004 vectors were selected on kanamycin plates and validated by bacterial colony PCR, followed by transformation into EHA105 by electroporation at 2500 V for about 5.5 ms. A single colony containing each target construct was confirmed by PCR and then used for genetic transformation of tobacco. EHA105-pCB2004-CsMYB6A and EHA105-pCB2004 were prepared for transformation. The leaf disc approach was used for tobacco transformation with 25 mg/L phosphinothricin selection.

### 2.7 Expression and purification of recombinant CsUGT72AM1

The cDNA of *CsUGT72AM1* were subcloned into the expression vector pMAL-c2X (New England Biolabs, MA, USA). The cloned gene sequences were also confirmed by colony PCR. The pMAL-*CsUGT72AM1* expression vector sand empty vectors were transformed into *E. coli* Novablue (DE3) competent cells (Novagen, Schwalbach, Germany). Recombinant proteins were purified according to the manufacturer’s instructions (New England Biolabs, MA, USA). Recombinant enzyme assays were carried out as described by Cui et al (Cui *et al*., 2016). The K_m_ and V_max_ of CsUGT72AM1 was determined using 5 mM UDP-glucose (UDP-Glc) as the sugar donor and 1.5–200 μM of flavonols as acceptors (kaempferol and quercetin) in phosphate buffer (pH 7.5). Reaction samples lacking recombinant proteins were used as blank controls. Reactions were stopped by mixing the reaction solutions with 100% methanol. All the kinetic assays were incubated at 30 °C for 10 min and repeated in triplicate.

### 2.8 Bioinformatics and statistical analyses

Multiple sequence alignment was performed using ClustalX. Phylogenetic tree was constructed using protein sequences from several plant MYBs and UGTs by Neighbor-Joining distance analysis. Branches corresponding to partitions reproduced in less than 50% bootstrap replicates are collapsed. The evolutionary distances were computed using the p-distance method.

Data were presented as mean ± SD. Statistically significant differences between the groups were determined with Student’s t-test using SPSS software (SPSS, Chicago, IL, USA). P<0.05 was considered to be statistically significant.

## 3 Results

### 3.1 Comparison of leaf color and flavonoid concentrations of different tea varieties

As shown in Figure 1A, anthocyanins were notably accumulated in ‘Mooma 1’ leaves. The color of young leaves and stems of ‘Mooma 1’ was bright red, whereas the color of wild-type was yellowish green (Figure 1C). The sections of tea leaves were sliced via free-hand sectioning to avoid the loss of anthocyanins. As shown in Figure 1E and G, anthocyanins in the purple-leaf tea variety were mainly accumulated and abundant in the palisade mesophyll (Pa) and xylem (Xy) cells, which were devoid of upper epidermal cells.

**Figure 1.**
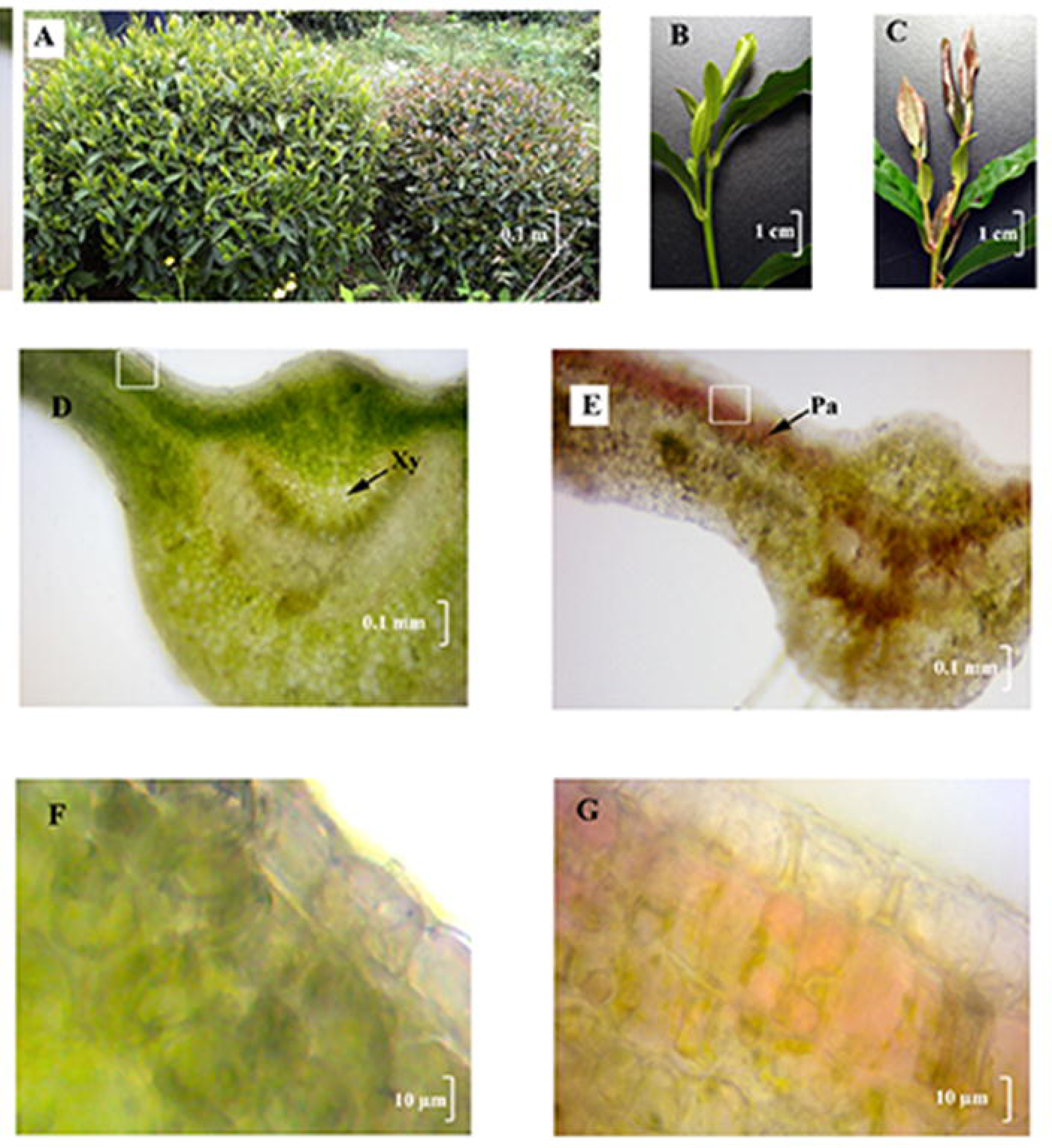
Phenotypic analysis of the red-leaf mutants of tea plants. (A) Wild-type tea plants and red-leaf mutants in the tea field; (B) and (C) Young branches of wild-type tea plants and red-leaf mutants; (D) and (F) Micromorphological images of the mutant leaf, × 40 fold and × 400 fold respectively; (E) and (G) Micromorphological images of the wild-type leaf staining with p-dimethylaminocinnamaldehyde (DMACA) - HCl, ×40 fold and ×400 fold respectively; Pa, palisade parenchyma; Gh, glandular hair; Xy, xylem.

### 3.2 Comparison of flavonoid compositions of different tea varieties

To investigate whether the flavonoid biosynthesis pathway of purple-leaf tea variety was different from the wild type, the flavonoid components of the fresh leaves were extracted and quantified. Catechins are the main compounds of flavonoids in the tea leaves, which approximately accounts for 13% of the dry weight. In summer, the amount of catechins in ‘Mooma 1’ leaves was significantly lower than the wild type (*P* < 0.05), whereas the amount showed no significant in spring (*P*=0.08).

The O-Glycosylated flavonol biosynthesis was strengthened in the purple-leaf tea variety. The total amounts of flavonols in ‘Mooma 1’ leaves (11.26±0.99 ng g^−1^ DW in spring and 17.52±1.06 ng g^−1^ DW in summer) were notably higher than in ‘Longjing 43’ leaves (9.91±0.38 ng g^−1^ DW in spring and 12.87±2.01 ng g^−1^ DW in summer), respectively. In addition, the amounts of flavonols in summer leaves were notably higher than in spring (*P*<0.05).

A representative HPLC profile of anthocyanins in ‘Mooma 1’ was presented in Figure 2 and Table 3. Eight anthocyanins were identified in ‘Mooma 1’ leaf using LC-MS analysis. Total amount of anthocyanins in the spring leaves (May 10th, average concentration: 591.87±51.1 ng g^−1^ DW) was notably higher than in the summer (July 20th, 451.11±19.02 ng g^−1^ DW, *P*<0.05), while anthocyanin concentration was very low in the wild type leaf (undetectable in two Seasons). In general, anthocyanin concentration in the purple-leaf tea was significantly higher than the wild-type cultivar, indicating that anthocyanin accumulation is responsible for the red coloration in tea.

**Figure 2.**
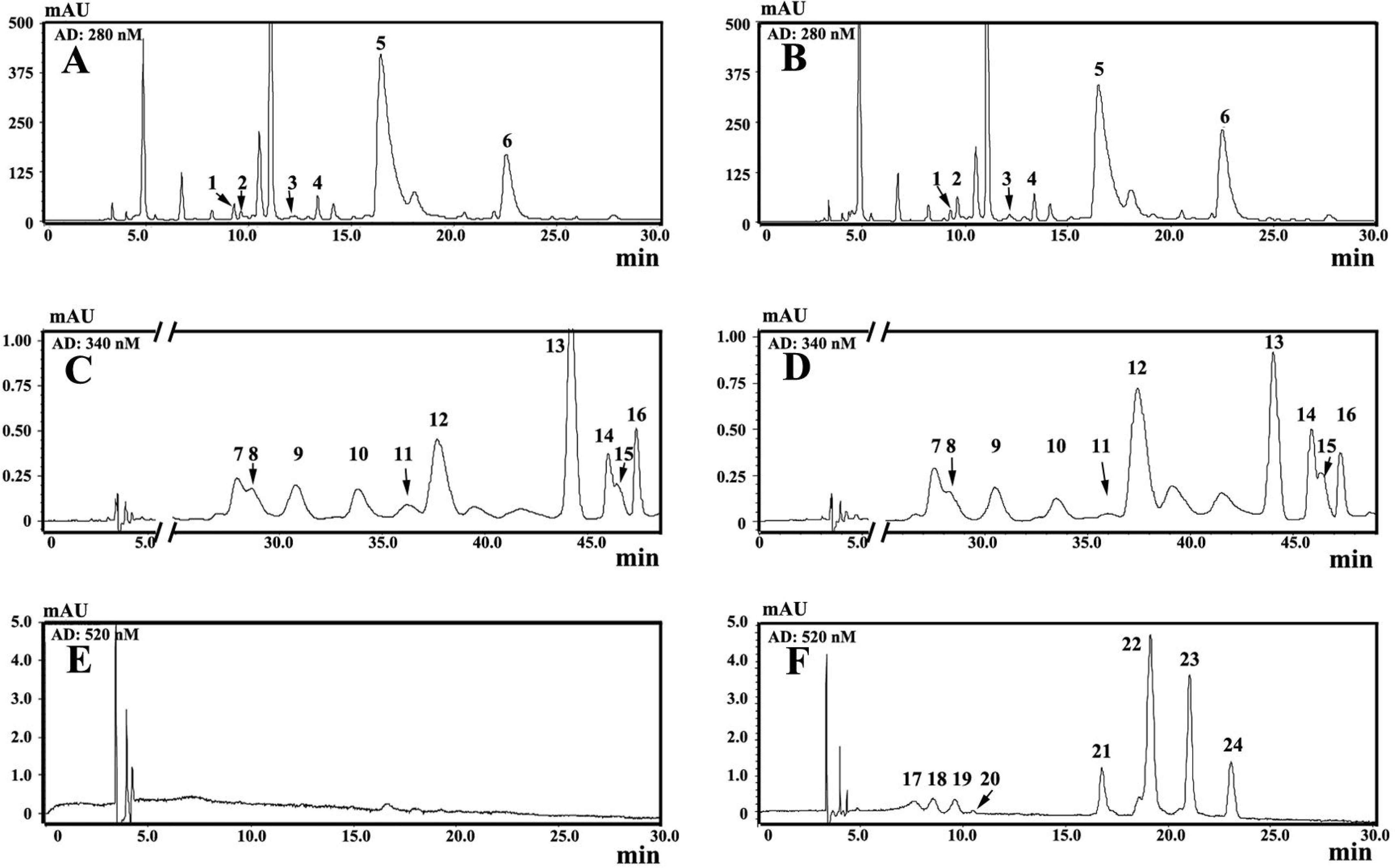
HPLC chromatogram of flavonoids in wild-type tea plants and red-leaf mutants. (A), (C), and (E) Representative HPLC chromatograms of flavonoids in wild-type tea plants at 280 nm, 345 nm, and 520 nm, respectively; (B), (D), and (F) Representative HPLC chromatograms of flavonoids in red-leaf mutantsat 280 nm, 345 nm, and 520 nm, respectively. Flavonoide compounds with peaks (from NO.1 to NO.24) were listed in Table 1.

### 3.3 Comparison of transcriptional difference between purple-leaf and wild-type tea

To deeply investigate anthocyanin and flavonol accumulation in purple-leaf tea, we carried out a comprehensive identification of transcriptional difference by RNA-Sequencing. Two normalized cDNA libraries from spring leaves of ‘Mooma1’ and wild type ‘Longjing43’ were sequenced. Following de novo assembly and redundancy reduction, we obtained a final set of 77,707 unigenes, with an average length of 1004 bp and N50 of 1675 bp (Table S2). Of these, 2,293 unigenes exhibited up-regulation, while 1,752 unigenes exhibited down-regulation in the purple-leaf transcript compared with the wild type (Figure S1). In total, 54,210 unigenes had hits in all five public databases with functional annotations (69.8% of the unigenes).

By mapping to the KEGG reference pathway, a total of 392 unigenes were assigned to the flavonoid biosynthesis pathway (including flavonoid, anthocyanin, and flavones and flavonol pathways). Among these annotated unigenes, whole sets of structural genes involved in the flavonoid biosynthesis pathway were indentified. All flavonoid genes were multiple genes, such as 5 *CsPAls*, 4 *CsDFRs*, 3 *CsCHSs*, and 3 *CsLARs* in tea plant (Figure 3). Unexpectedly, all the highly (FPKM > 100) and moderately (FPKM > 10) expressed genes, which are related to flavonoid biosynthesis, found no significant up-regulation in the purple variety. Only two low expressed genes, *CsF3’H* and *CsDFR2* (FPKM<10) showed 2- and 1.9-fold higher in purple-leaf than in green-leaf. Several CsUGTs encoding terminal enzymes by galactosylated modification of flavonoids were remarkably up-regulated in ‘Mooma1’ leaf. Especially, the expression of CsUGT72AM1 and CsUGT3 in ‘Mooma1’ leaf was increased by 4.2- and 2.5-folds respectively, compared with the value in wild-type tea. The result was further confirmed by qRT-PCR.

**Figure 3.**
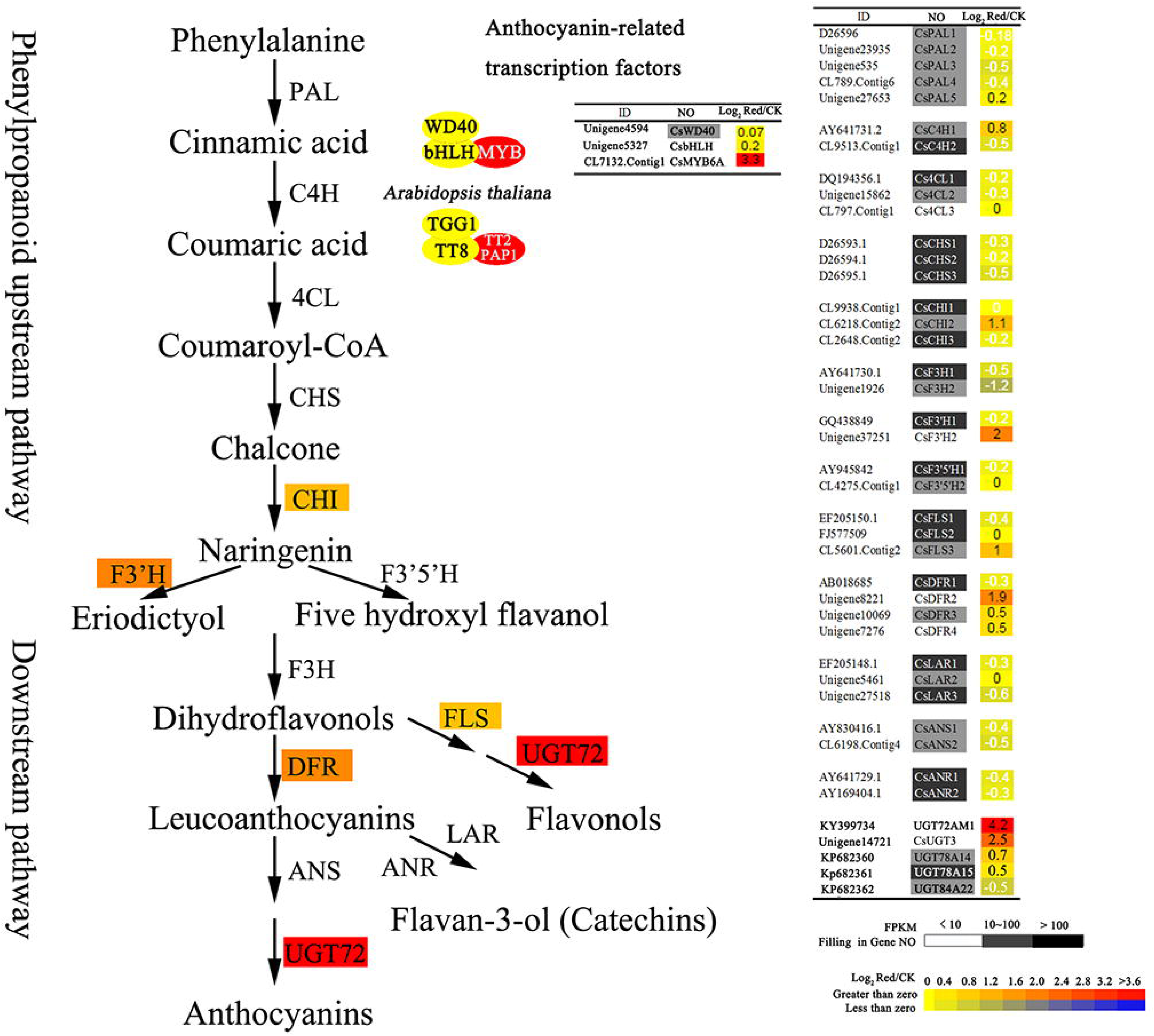
Comparison of transcripts coding flavonoid biosynthetic enzymes in leaves of purple-leaf mutants and wild-type tea plants. Transcripts coding enzymes involved in the flavonoid pathway were identified by screening the Camellia sinensis RNA-sequence libraries and NCBI database (www.ncbi.nlm.nih.gov). Black, gray, and white squares enveloping the gene No. represents transcript level are high (FPKM > 100), medium (100 > FPKM > 10), and low (FPKM<10), respectively. Gradient color square, from yellow to red, on the data of Log_2_^Red/CK^ representing an increasing gene expression level from 0 to > 3 fold in the purple-leaf mutants was compared with the wild-type green tea plant.

MYB-bHLH-WD40 (MBW) complexes, activating anthocyanin biosynthesis, were also investigated with the transcriptome of purple-leaf. The homologues of *AtMYB113* (*CsMYB6A*, NO. CL7132), *AtTT8* (*CsTT8*, NO. Unigene5327) and *AtTTG1* (*CsTTG1*, NO. Unigene4594) were identified. However, *CsTT8* and *CsTTG1* expression levels were not significantly up-regulated in ‘Mooma1’ leaf. Only the relative expression level of *CsMYB6A* was significantly up-regulated in the leaves of ‘Mooma1’, and was increased by 3.3-folds compared with wild-type.

126 R2R3-MYBs from *Arabidopsis thaliana* and CsMYB6A were used in phylogenetic tree construction (Figure 4). Phylogenetic analysis indicated CsMYB6A, attached to subgroup 6 (Sg6), was most similar to AtMYB113 and shares 63% identity (Figure 4). MYBs in Sg6 are reported regulators of anthocyanin accumulation (Hichri *et al*., 2011). CsMYB6A-GFP fusion transient expression showed that CsMYB6A protein was exclusively localized in the nucleus (Figure S2).

**Figure 4.**
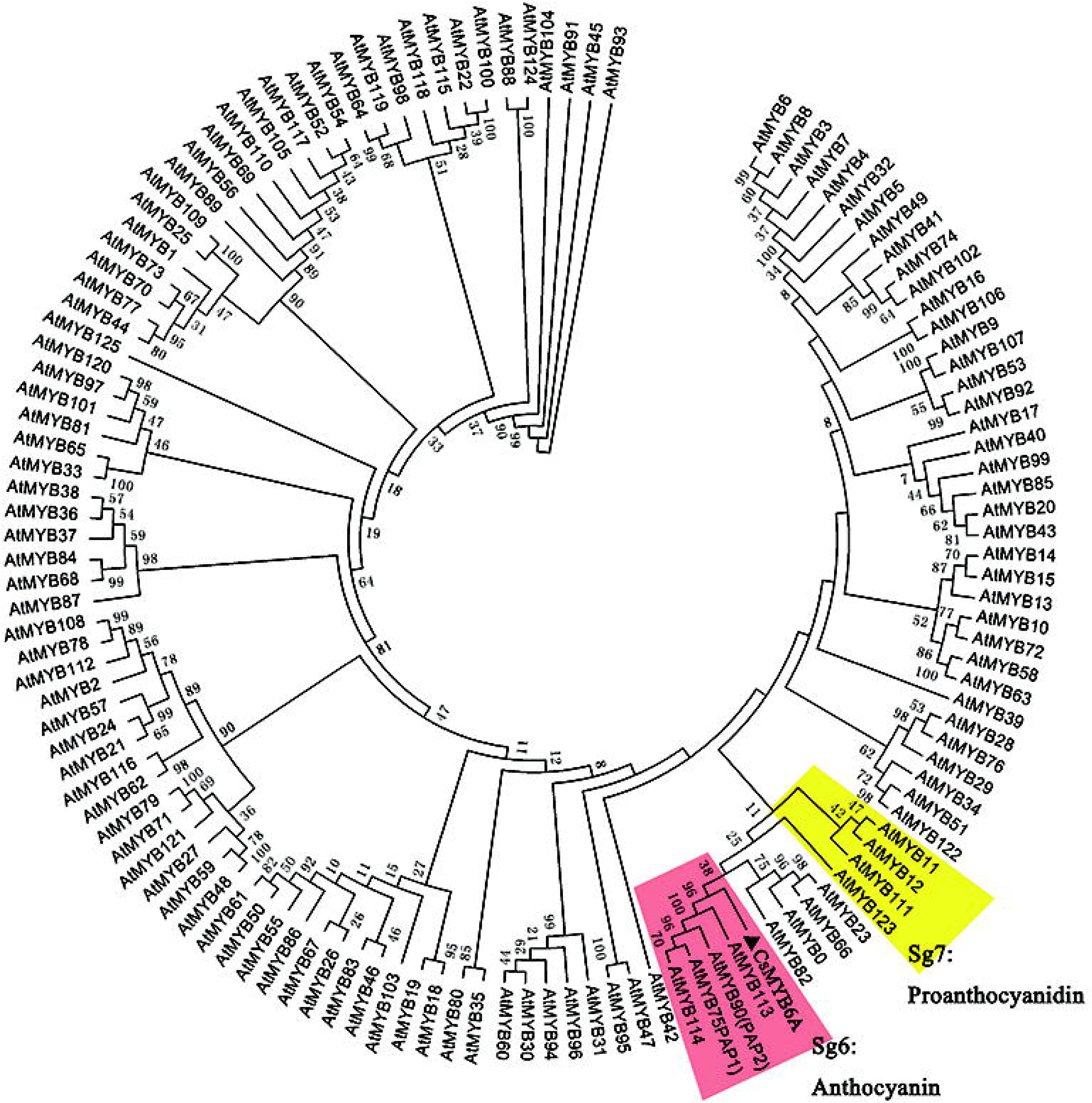
Phylogenetic tree of R2R3-MYB transcription factors. The protein sequences contained CsMYB6A from Camellia sinensis and AtMYBs from Arabidopsis thaliana. The phylogenetic tree was constructed using MEGA 5 with 1000 bootstrap replicates. Numbers indicate the percentage of consensus support.

### 3.4 Functional analysis of the *CsMYB6A* gene in *Nicotiana tabacum*

A R2R3-MYB (PAP1) of *Arabidopsis thaliana* could enhance the accumulation of anthocyanins inducing the purple color of the leaf and stem (Borevitz *et al*., 2000). The 35S:CsMYB6A, 35S:AtPAP1 (as positive control), and empty plasmid (as negative control) vectors were introduced into Tobacco ‘G28’ (*Nicotiana tabacum* ‘G28’). About 10 independent transgenic tobacco plants of different genes were obtained. The leaves of *CsMYB6A* or *AtPAP1* transgenic plants exhibited a clear color change from green of the control host to purple (Figure 5A). The concentrations of anthocyanins in the leaf of *CsMYB6A* and *AtPAP1* transgenic tobacco, as detected by HLPC method, were 240 and 340 ng g^−1^DW respectively, while undetected in the leaf of empty vector control (Figure 5B). Additionally, the accumulation of flavonols in the leaf of *CsMYB6A* or *AtPAP1* transgenic tobacco was remarkably enhanced by 1.89 and 4.15-folds respectively, compared to the vector control. Environmental temperature and light are considered as important factors that affect anthocyanin biosynthesis in garden plants. Our data also confirmed that low temperature and red light accelerated the anthocyanin accumulation in the transgenic tobacco lines (Figure S3).

**Figure 5.**
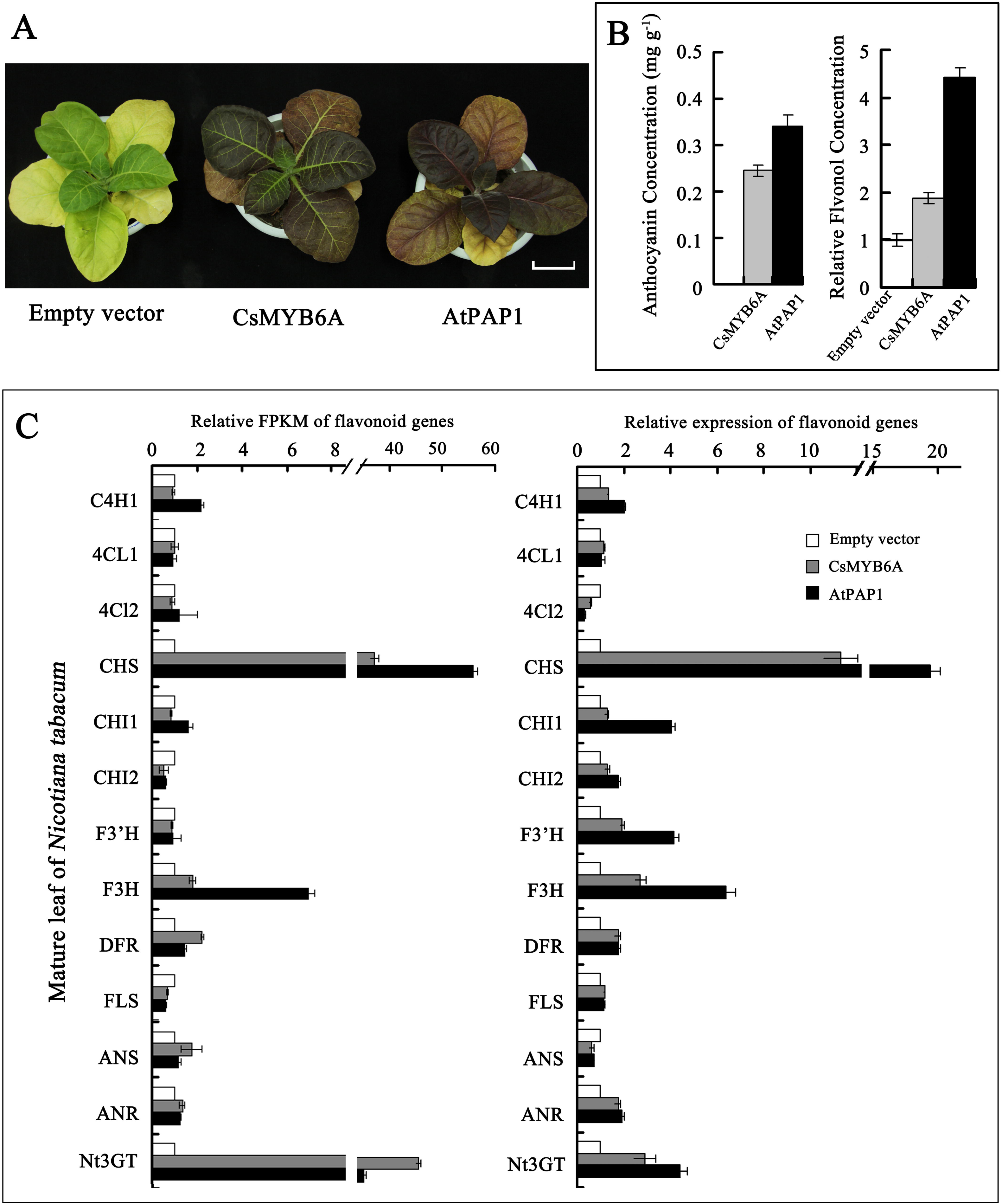
Phenotypic analysis of transgenic tobacco overexpressing *CsMYB6A* and *AtPAP1*. (A) Leaves of transgenic plants (CsMYB6A and AtPAP1) in comparion with empty-vector transgenic plants; (B) Total concentrations of anthocyanins and flavonols of transgenic plants. The relative flavonol concentration was calculated as the ratio between the total peak area at 350 nm; (C)The relative gene expression involved in flavonoid biosynthesis in transgemic plants compared with the empty vector. Relative FPKM and expression data were obtained from RNA-sequencing and qRT-PCR analysis, respectively.

To investigate whether the flavonoid biosynthesis pathway was affected by over-expression of *R2R3-MYBs*, the flavonoid pathway genes, including *C4H*, *4CL*, *CHS*, *CHI*, *F3’H*, *F3H*, *DFR*, *FLS*, *LAR*, *ANS*, *ANR*, and *3GT* were examined by qRT-PCR in vector control and transgenic lines with β-actin (accession number: EU938079) as reference gene (Figure 5C). The expression levels of *F3H*, and *3GT* genes in the mature leaf of transgenic lines was significantly increased by 2-folds in comparison to the vector control lines, especially the level of *CHS* was enhanced by 10-folds as compared to the control lines. Therefore, the expression of these genes was stimulated by the over-expression of CsMYB6A or AtPAP1 in transgenic lines. The result was also confirmed by qRT-PCR (Figure 5C).

### 3.5 Heterologous expression and enzymic analysis of the recombinant *CsUGT72AM1*

RNA-Sequencing data showed that several CsUGTs, especially *CsUGT72AM1* and *CsUGT3*, were remarkably up-regulated in ‘Mooma1’ leaf, compared with the value in wild-type tea. Our previous research had identified a UDP-glycosyltransferase, *CsUGT78A14*, which is responsible for the biosynthesis of flavonol 3-*O*-glucosides (Cui *et al*., 2016). The expression of *CsUGT78A14*was unremarkably up-regulated in the ‘Mooma1’ leaf compared with wild-type control. Phylogenetic analysis, referring to *Arabidopsis* UGTs, indicated CsUGT72AM1, CsUGT3 and CsUGT78A14 are the most similar to AtUGT72E1, AtUGT91A1, and Nt3GT, respectively (Figure 6).

**Figure 6.**
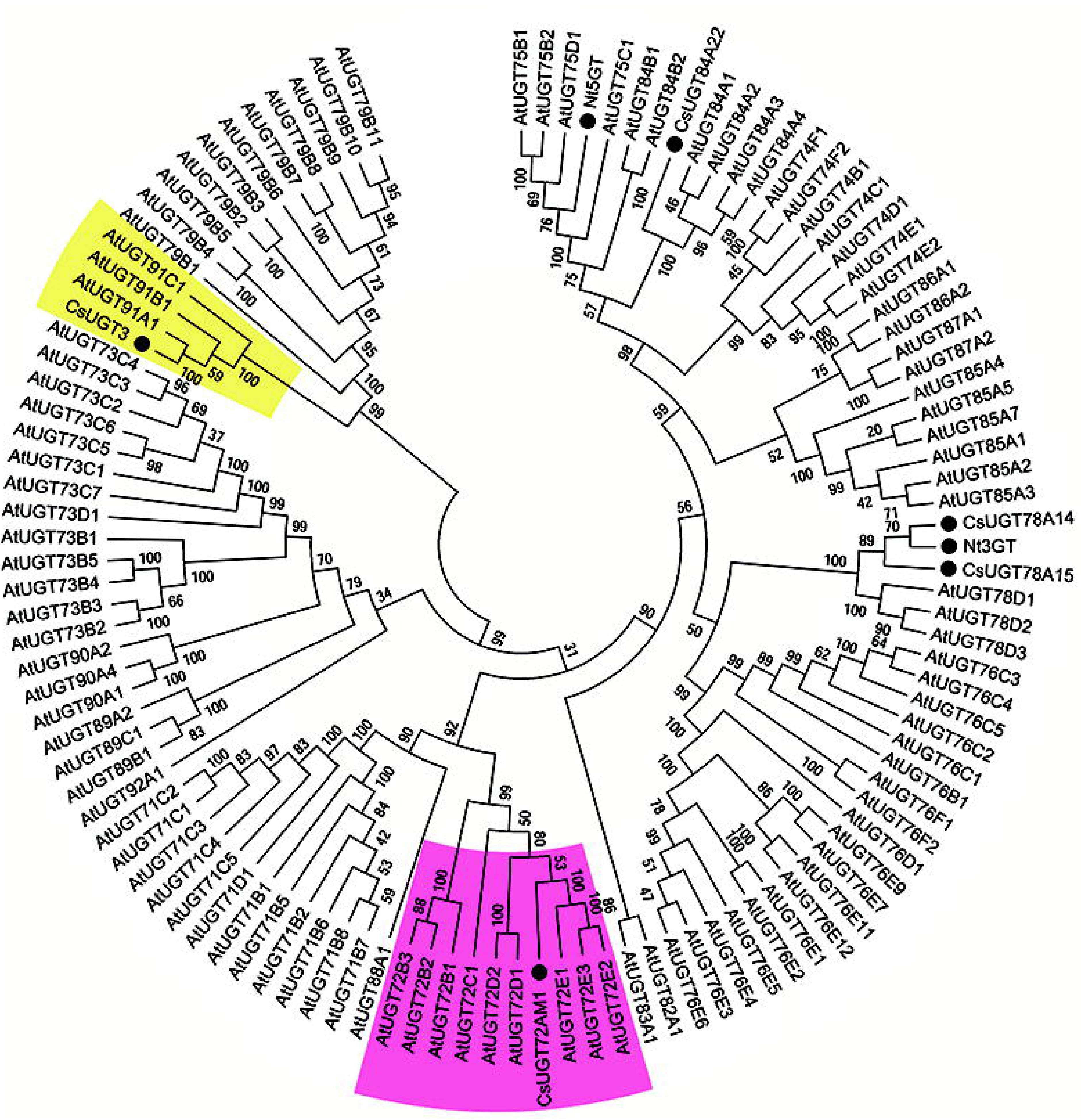
Phylogenetic tree of UGT proteins. The protein sequences contained CsUGT72AM1, CsUGT78A14, CsUGT3 from *Camellia sinensis* and AtUGTs from *Arabidopsis thaliana*. The phylogenetic tree was constructed using MEGA 5 with 1000 bootstrap replicates. Number indicates the percentage of consensus support.

The enzymatic characteristics of CsUGT72AM1 and CsUGT78A14 proteins that were expressed *Escherichia coli* cells were evaluated. The purified recombinant CsUGT72AM1 proteins, in accordance with rCsUGT78A14, can catalyze the glycosylation of quercetin or cyanidin at 3-OH group with UDP-Glucose as the sugar donors in vitro (Figure 7A, B). However, the recombinant enzyme demonstrated low activity for anthocyanins, and exhibited remarkable substrate inhibition, which was consistent with previous published reports (Kovinich *et al*., 2010).

**Figure 7.**
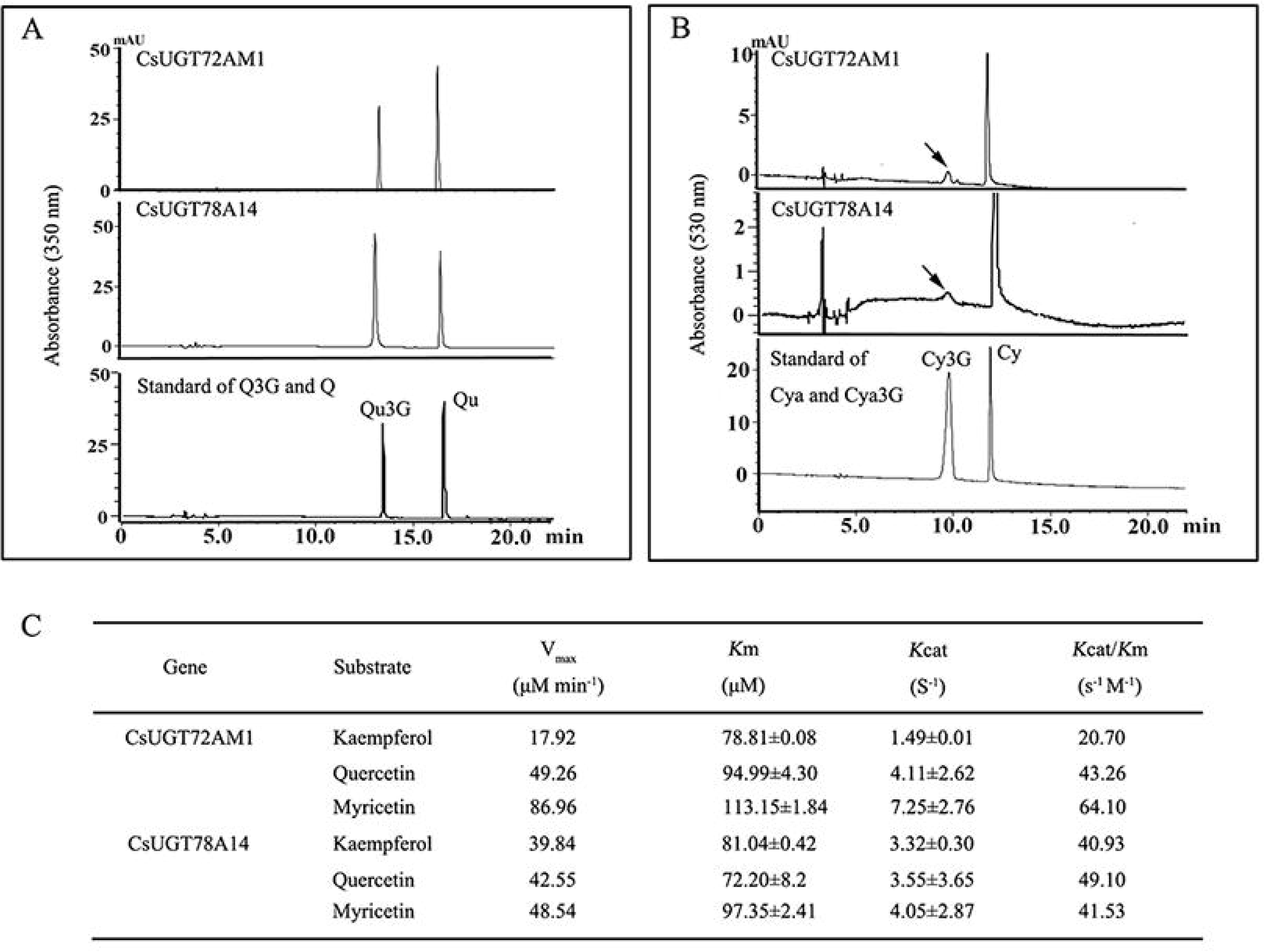
Comparison of the enzymatic activities of the CsUGT72AM1 with CsUGT78A14. (A) HPLC chromatograms for the enzymatic product of the recombination proteins with quercetin as flavonol acceptor. (B) HPLC chromatograms for the enzymatic product of the recombination proteins with cyanidin as anthocyanin acceptor. (C) Kinetic parameters of the recombinant proteins for quercetin.

The kinetic parameters of these two recombinant glucosyltransferases for flavonols (kaempferol, quercetin, and myricetin) were determined in phosphate buffer at pH 7.5. The recombinant enzymes demonstrated highest activity for flavonols. The Km values of UGT72AM1 for kaempferol, quercetin, and myricetin were 71.81, 94.99, and 113.15 μM with Vmax values of 17.92, 49.26, and 86.96 nmol min^−1^, respectively. For UGT78A14, the Km and Vmax values of kaempferol, quercetin, and myricetin were 81.04, 72.20, and 97.35 μM and 39.84, 42.55, and 48.54 nmol min^−1^, respectively (Figure 7C).

## 4. Discussion

### 4.1 Flavonoid components in the leaves of purple-leaf tea

Several health benefits were observed with tea products, which contain high concentrations of flavonoid components. This drove the scholar’s initiatives to explore the specialty of tea varieties, including white tea, yellow tea, anthocyanin-rich tea. Several purple-leaf tea varieties were reported in different tea planting areas (Joshi *et al*., 2015; Kerio *et al*., 2013; Lv *et al*., 2015). A new purple-leaf tea variety ‘Mooma 1’ was reported in this paper, which was selected from the natural hybrid population of ‘Longjing 43’. The buds and young leaves of ‘Mooma 1’ displayed bright red color, where the anthocyanins were remarkably accumulated in the palisade mesophyll tissue, but were absent in the epidermis. A previous study demonstrated that the putative function of antioxidative defense in the leaves was more possible for anthocyanins located in the mesophyll than in the epidermal vacuoles (Kytridis and Manetas, 2006).

Flavonoid compounds are considered to be the most important quality parameters of tea products due to their impact on color and taste properties. Catechins are responsible for astringency and bitterness (Kallithraka et al., 1997). Flavonols induce a velvety and mouth-coating sensation at very low concentration (Scharbert *et al*., 2004). Anthocyanins add to the briskness by complexing with catechins and theaflavins (Joshi *et al*., 2015). Purple-leaf tea products show more astringency with better mouth feel and sweet after taste than green-leaf tea. Average concentrations of total anthocyanins in the spring or summer leaves were 591.87±51.1 and 451.11±19.02 ng g^−1^ DW, respectively, which was very low in the wild type leaves. Additionally, the total amounts of O-Glycosylated flavonols in ‘Mooma 1’ leaves were significantly higher than in the control leaves.

The biosynthetic pathways of anthocyanins, flavonols, and flavan-3-ols (catechins) share common steps in the phenylpropanoid and flavonoid pathways (from PAL to F3H). Each class of flavonoid was synthesized by a multienzymatic step reaction branching from the common flavonoid pathway (Figure 3). Several studies have shown the competitive relationship between different flavonoids due to competition for the substrate. Interestingly, our results indicated that anthocyanins and flavonols have a synergistic relationship, and both showed significant increase in the purple-leaf variety compared with the control variety. Both the compounds may have competition with catechins in purple-leaf variety, because the total amount of catechins in purple leaves was slightly decreased compared with the control. However, further research is required to uncover the association between anthocyanin and flavonol accumulation.

### 4.2 An R2R3-MYB transcription factor, CsMYB6A, promotes flavonoid accumulation in purple-leaf tea

Several R2R3-MYB transcription factors, combined with other transcription factors (bHLH and WD40), are known to be involved in the regulation of flavonoid biosynthesis. *AN2* from *Perunia hybrid* (Quattrocchio et al., 1993), *C1* from *Zea mays* (Consonni et al., 1993), and *PAP1* and *PAP2* from *Arabidopsis thaliana* (Borevitz et al., 2000) all encode MYB proteins. These regulate the accumulation of different anthocyanin pigments. Pattanaik confirmed that anthocyanin related MYB interacts with other heterologous species, *bHLH,* to activate the expression of key flavonoid pathway genes, including *CHS* and *DFR*, which induces anthocyanin synthesis (Pattanaik *et al*., 2010). These results indicate that regulatory anthocyanin genes are conserved between species.

Our previous study predicted R2R3-MYB genes in flavonoid biosynthesis of *C. sinensis*, while Sg6 CsMYB genes were lacked in the wild-type green-leaf tea, (Zhao *et al*., 2013). In the current study of ‘Mooma 1’ purple-leaf cultivars, we found that the expression of the CsMYB6A was most similar to AtMYB113, and was consistent in regulating anthocyanin synthesis. The different expression levels of CsMYB6A in different cultivars might be due to the transcriptional modification or post-transcriptional modification (Li *et al*., 2012; Sun *et al*., 2016). Furthermore, we also investigated the over-expression of CsMYB6A and AtPAP1 genes in transgenic tobacco, G28. Results showed that there was significant up-regulation of structural flavonoid genes, especially CHS and anthocyanin 3-O-glucosyltransferase (A3T) in both CsMYB6A and AtPAP1 transgenic tobacco lines compared with empty-vector control. Interestingly, the expression levels of CsUGT78A14, the homologous gene of 3GT, demonstrated no significant up-regulation in the leaves of ‘Mooma 1’, while several other UGTs, especially CsUGT72AM1, were obviously up-regulated compared with green-leaf tea. These results indicated that the divergent target genes were responsible for the species-specific differences in regulatory networks.

### 4.3 CsUGT72AM1, an UGT gene, catalyzes the glycosylation of flavonoids

UDP-glycosyltransferases are the final enzymes produced in anthocyanin and flavonol biosynthesis. 78 UGT family genes, such as *UGT78G1* (Modolo *et al*., 2009), *UGT78D2* (Tohge *et al*., 2005) were activated with anthocyanidins, flavonols, flavones, coumestans, pterocarpans, and isoflavones, and were involved in the O-glucosylation of anthocyanins. In our previous study, CsUGT78A14 was found to be responsible for the biosynthesis of flavonol 3-O-glucosides (Cui *et al*., 2016).

A novel UGT gene, CsUGT72AM1, showed higher expression level in ‘Mooma1’, and was screened and cloned from purple-tea leaves. Phylogenetic study indicated that CsUGT72AM1, which is similar to AtUGT72E1, was clustered into the 72 UGT family of glucosyltransferases. The 72 UGT subgroup, studied in several previous studies, was able to catalyze the formation of monolignol 4-O-glucose that is involved in the biosynthesis of lignin (Yonekura-Sakakibara and Hanada, 2011). Several UGT family genes can catalyze the O-glucosylation of flavonoids. For example, UGT72L1 from *Medicago truncatula*, which was regulated by *Arabidopsis* R2R3-MYB (TT2) was specifically active towards the epicatechin, formatting epicatechin 3’-O-glucoside (Pang *et al*., 2008).

In this paper, we compared the enzymatic characteristics of CsUGT72AM1 and CsUGT78A14 through *in vitro* assays. Our enzymatic assays confirmed that both CsUGT72AM1 and CsUGT78A14 demonstrated catalytic activity as a flavonol or anthocyanin 3-O-glucosyltransferase. The recombinant enzymes CsUGT72AM1 and CsUGT78A14 were highly catalyzed by the addition of glycosyl group from UTP-glucose to flavonols. CsUGT72AM1 displayed broad substrate specificity, by in vitro experiments, recognizing the flavonoid substrates, including naringenin (N), Eriodictyol (E), kaempferol (K), quercetin (Q), and myricetin (M), as acceptor molecules (Data not shown in this paper). These results indicated that both CsUGT72AM1 and CsUGT78A14 genes are likely to be involved in the biosynthesis of flavonoid 3-O-glycoside compounds in tea plants.

Taken together, our research acquired the key genes, including CsMYB6A and CsUGT72AM1, which regulates the accumulation of anthocyanin and flavonol in purple-leaf tea. CsMYB6A transgenic results indicated that the regulatory mechanism of flavonoid pathway was conserved between species (Quattrocchio *et al*., 1998). In future, we would focus on exploiting MYB-target genes using interaction methods. Therefore, CsMYB6A and related CsUGTs are the first characterized genes that are involved in the anthocyanin biosynthesis in tea leaf organs. Their identification provides advances in understanding the flavonoid biosynthesis in non-alcoholic beverage crops.

---

**Figure S1 Summary of differentially expressed unigenes in red-leaf mutants and wild-type tea plants.** Red, Green and blue spots represent up-regulated, down-regulated, no-change unigenes, respectively.

**Figure S2 Subcelleular localization of CsMYB6A protein** (A) Red fluorescence of chloroplast; (B) Green fluorescence of 35S:CsMYB6A-GFP fusion protein; (C) Merged slice of (A) and (B); (D), (E) and (F) highly magnificent slice of white square of (A), (B), and (C).

**Figure S3 Effect of temperature and illumination on anthocyanin accumulation in transgenic tobacco overexpressing *CsMYB6A*.** (A) Phenotype of transgenic plants in different conditions; (B) Comparison of total anthocyanin concentrations of transgenic plants in different conditions. (C)Comparison of leaf color of transgenic plants in different illumination conditions.

